# Cotranslational folding of a periplasmic protein domain in *Escherichia coli*

**DOI:** 10.1101/2021.02.06.430026

**Authors:** Hena Sandhu, Rickard Hedman, Florian Cymer, Renuka Kudva, Nurzian Ismail, Gunnar von Heijne

## Abstract

In Gram-negative bacteria, periplasmic domains in inner membrane proteins are cotranslationally translocated across the inner membrane through the SecYEG translocon. To what degree such domains also start to fold cotranslationally is generally difficult to determine using currently available methods. Here, we apply Force Profile Analysis (FPA) – a method where a translational arrest peptide is used to detect folding-induced forces acting on the nascent polypeptide – to follow the cotranslational translocation and folding of the large periplasmic domain of the *E. coli* inner membrane protease LepB *in vivo*. Membrane insertion of LepB’s two N-terminal transmembrane helices is initiated when their respective N-terminal ends reach 45-50 residues away from the peptidyl transferase center (PTC) in the ribosome. The main folding transition in the periplasmic domain involves all but the ~15 most C-terminal residues of the protein and happens when the C-terminal end of the folded part is ~70 residues away from the PTC; a smaller putative folding intermediate is also detected. This implies that wildtype LepB folds post-translationally *in vivo*, and shows that FPA can be used to study both co- and post-translational protein folding in the periplasm.

## Introduction

Most secreted proteins are translocated through Sec-type translocons in the bacterial inner membrane or the eukaryotic endoplasmic reticulum membrane in an unfolded state, and fold once they emerge on the *trans* side of the membrane [1]. The folding of proteins that are cotranslationally translocated across the membrane can be followed by tracking the formation of specific disulfide bonds or by assaying the appearance of enzymatically active domains as the protein emerges from the bacterial SecYEG or the eukaryotic Sec61 translocation channel [2–5], but this provides only a gross indication of the formation of folded structure.

In contrast, the cotranslational folding of cytosolic proteins can be studied in considerable detail, both *in vitro* and *in vivo*, using FRET-based assays [6, 7], protease-protection assays [6, 8–13], NMR [14–19], cryo-electron microscopy [20–22], optical tweezer experiments [23–26], and force-profile analysis (FPA) [7, 13, 20–22, 26–31]. These methods can track where in the ribosome exit tunnel a protein starts to fold, whether any folding intermediates form before the nascent protein has been fully synthesized, and to what extent the presence of the ribosome affects the folding kinetics and the thermodynamic stability of the folded state.

Among these methods, FPA is unique in that it can be applied both *in vitro* and *in vivo*, and because it can be used to analyze any kind of event that results in a pulling force on the nascent polypeptide chain, including protein folding. Here, we show that FPA can be used to follow the cotranslational folding of the ~250 residues periplasmic domain in the *E. coli* inner membrane protein LepB, and show that the ~15 C-terminal residues of LepB are not involved in the main folding transition.

## Results

### The force profile analysis (FPA) assay

Translational arrest peptides (APs) are short stretches of polypeptide that interact with the ribosome exit tunnel to pause translation when the last codon in the portion of the mRNA that codes for the AP is located in the ribosomal A-site [32]. The stalling efficiency of APs is sensitive to pulling forces acting on the nascent chain [26, 33, 34], and APs can therefore be used as force sensors to report on cotranslational events that generate pulling forces, such as protein folding or insertion of transmembrane segments into the membrane.

In FPA, a force-generating domain in a polypeptide is placed at increasing distances upstream of an AP, and the degree of translational stalling is measured for the corresponding series of protein constructs. A plot of the stalling efficiency *vs*. chain length – a force profile (FP) – provides a map of cotranslational force-generating events with up to single-residue resolution. Further, by using APs of different stalling strengths, FPs can be fine-tuned to optimally reflect different kinds of force-generating events [35, 36].

Here, we explore the possibility to use FPA to follow the cotranslational folding of a periplasmic domain in an *E. coli* inner membrane protein as it emerges from the SecYEG translocon into the periplasm. As illustrated in Fig. 1a, in constructs where the nascent chain is long enough that it can be stretched to the point that a protein domain (or a part of a domain that can form a stable folding intermediate) can reach sufficiently far into the periplasm to start to fold, some of the free energy gained upon folding will be converted to tension in the nascent chain, generating a pulling force on the AP and thereby reducing the stalling efficiency. As shown below, we find that protein folding in the periplasm is readily amenable to analysis by FPA.

**Figure 1.**
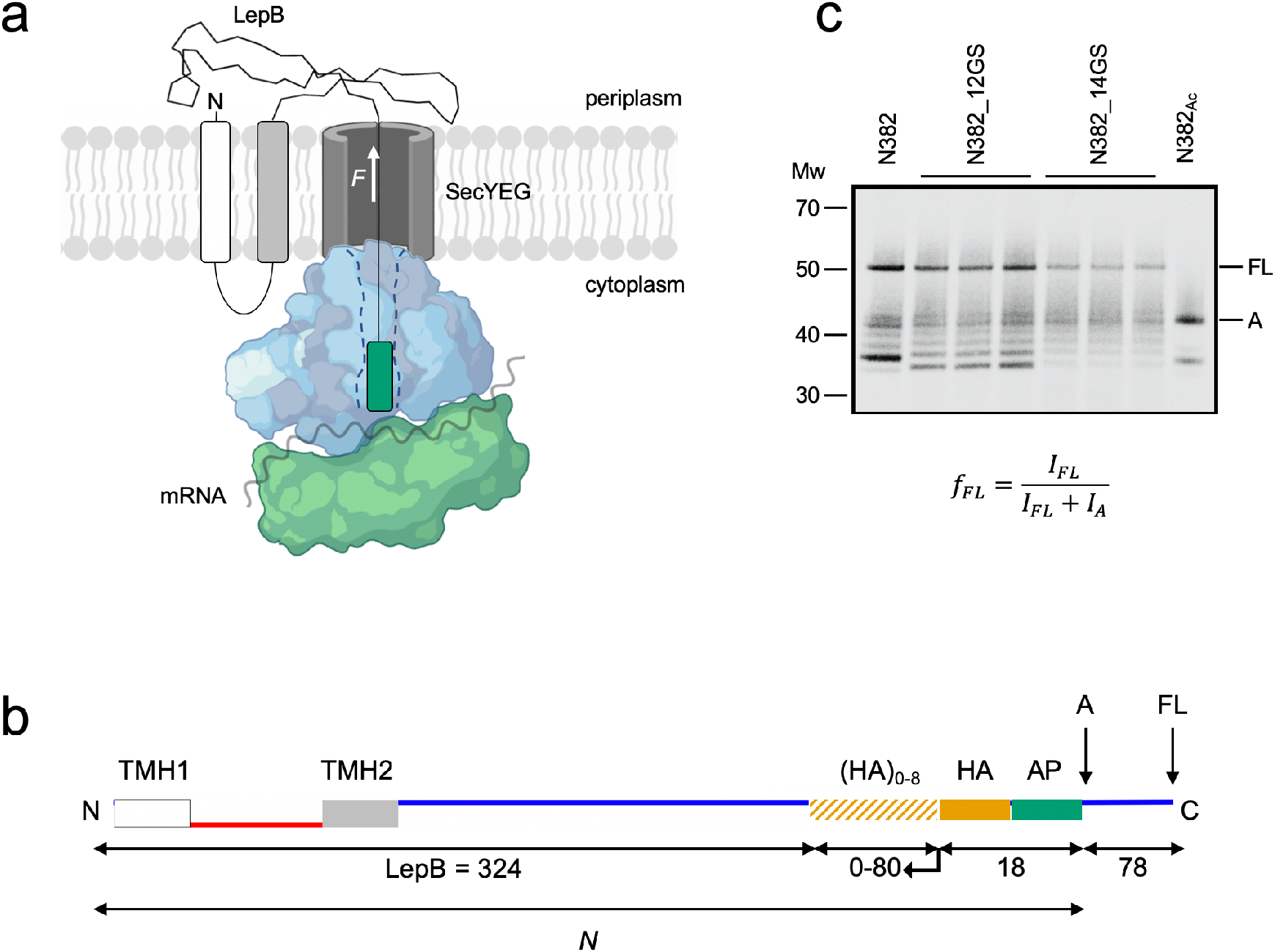
Force profile analysis of cotranslational protein folding in the periplasm. (a) Schematic representation of a cotranslationally arrested ribosome carrying a partially translated LepB nascent chain. Folding of the LepB periplasmic domain generates a force *F* on the nascent chain that is transmitted to the AP (in green), leading to less efficient stalling and increased production of full-length LepB. (b) Construct design. HA tags, AP, and C-terminal tail are indicated, together with the C-terminal ends of the arrested (*A*) and full-length (*FL*) products. The longest construct (*N* = 422) has 9 HA tags fused to the C terminus of LepB; shorter constructs are made by C-to-N-terminal truncations starting between the 8^th^ and 9^th^ HA tag, as indicated by the arrow. Cytoplasmic (red) and periplasmic (blue) parts are indicated. The 18-residue HA+AP segment and the C-terminal tail are constant in all constructs. (c) Representative SDS-PAGE gel for quantification of fraction full-length protein (*f*_*FL*_). Lane 1 is construct *N* = 382, lanes 2-4 a triplicate of construct *N* = 382 with the last 24 residues of LepB replaced by 12 GS-repeats, lanes 5-7 construct *N* = 382 with the last 28 residues of LepB replaced by 14 GS-repeats, and lane 8 is an arrest control (*Ac*) for construct *N* = 382 with a stop codon in place of the last Pro residue in the AP. The ladder of bands below the arrested (*A*) form of the protein likely results from ribosome stacking behind the arrested ribosome.

### Cotranslational membrane insertion and folding of LepB

*E. coli* LepB is a 324-residue inner membrane protease. It is anchored in the inner membrane by two N-terminal transmembrane helices (TMH1, TMH2) and has a large C-terminal periplasmic domain that contains the active site [37, 38]. The signal recognition particle interacts with TMH1 as it emerges from the ribosome exit tunnel at a nascent chain length of ~45 residues [39–41], targeting the ribosome-nascent chain complex (RNC) to the SecYEG translocon. TMH1 then inserts into the inner membrane in the N_out_-C_in_ orientation with its N-terminus facing the periplasm. Once TMH2 emerges, it inserts into the membrane in the opposite N_in_-C_out_ orientation and initiates cotranslational, SecA-dependent translocation of the C-terminal periplasmic domain through the SecYEG translocon [42].

To follow the cotranslational insertion of the transmembrane helices and the folding of the periplasmic domain, we made a series of constructs composed of progressively longer N-terminal parts of the LepB protein followed by a 9-residue HA-tag, a 8-residue SecM(*Ms*) AP (a relatively strong AP [33]), and a 78-residue C-terminal tail, Fig. 1b (see Supplementary Table 1 for sequences of all constructs). We also made longer constructs where full-length LepB is followed by linkers composed of additional HA-tags, the SecM(*Ms*) AP, and the C-terminal tail.

All constructs were expressed in *E. coli* MC1061 and pulse-labelled by [^35^S] Met for 3 min. LepB products were immunoprecipitated by an HA antiserum, separated by SDS-PAGE, and visualized by phosphorimaging. As seen in the example given in Fig. 1c, two main LepB species are produced, corresponding to arrested nascent chains (*A*) and full-length protein including the C-terminal tail (*FL*). The *A* and *FL* bands were quantitated and the fraction full-length protein was calculated as *f*_*FL*_ = *I_FL_*/(*I_FL_*+*I_A_*), where *I_A_* and *I_FL_* are the intensities of the *A* and *FL* protein bands, respectively.

A force-profile (FP) plot of the *f*_*FL*_ values for the different LepB constructs versus the length *N* of the construct (*N* = the number of residues counting from the N terminus of LepB to the critical proline residue at the C-terminal end of the AP) is shown in Fig. 2a. Five peaks (I-V) are seen in the FP. Peak I reaches half-maximal amplitude at *N*_*start*_ = 53 residues and has its maximum at *N*_*max*_ = 55 residues. The hydrophobic section of TMH1 encompasses residues 4-22, hence the N-terminal end of TMH1 is 50 residues away from the polypeptide transferase center (PTC) in the *N* = 53 construct, Fig. 2b. In a previous study, using the SecM(*Ms*) AP [33] we found that the main peak in a force profile representing the insertion of an artificial TMH of composition 6L/13A into the *E. coli* inner membrane reaches half-maximal amplitude when the N-terminal end of the TMH is ~50 residues away from the PTC, suggesting that peak I is generated by the insertion of LepB TMH1 into the inner membrane. A double substitution [F^5^, L^7^ → R^5^, R^7^] that makes the N-terminus of TMH1 less hydrophobic results in a reduced *f*_*FL*_ value at *N* = 55 (green data point), as expected. The assignment of TMH1 to peak I is further corroborated by previous crosslinking studies [43, 44] and by a recent continuous-translation *in vitro* study in which a FRET signal between an acceptor attached to the N-terminal Met residue of LepB and a donor placed at the cytoplasmic entry to the SecYEG channel is seen when the LepB nascent chain reaches a length of ~50 residues [45].

**Figure 2.**
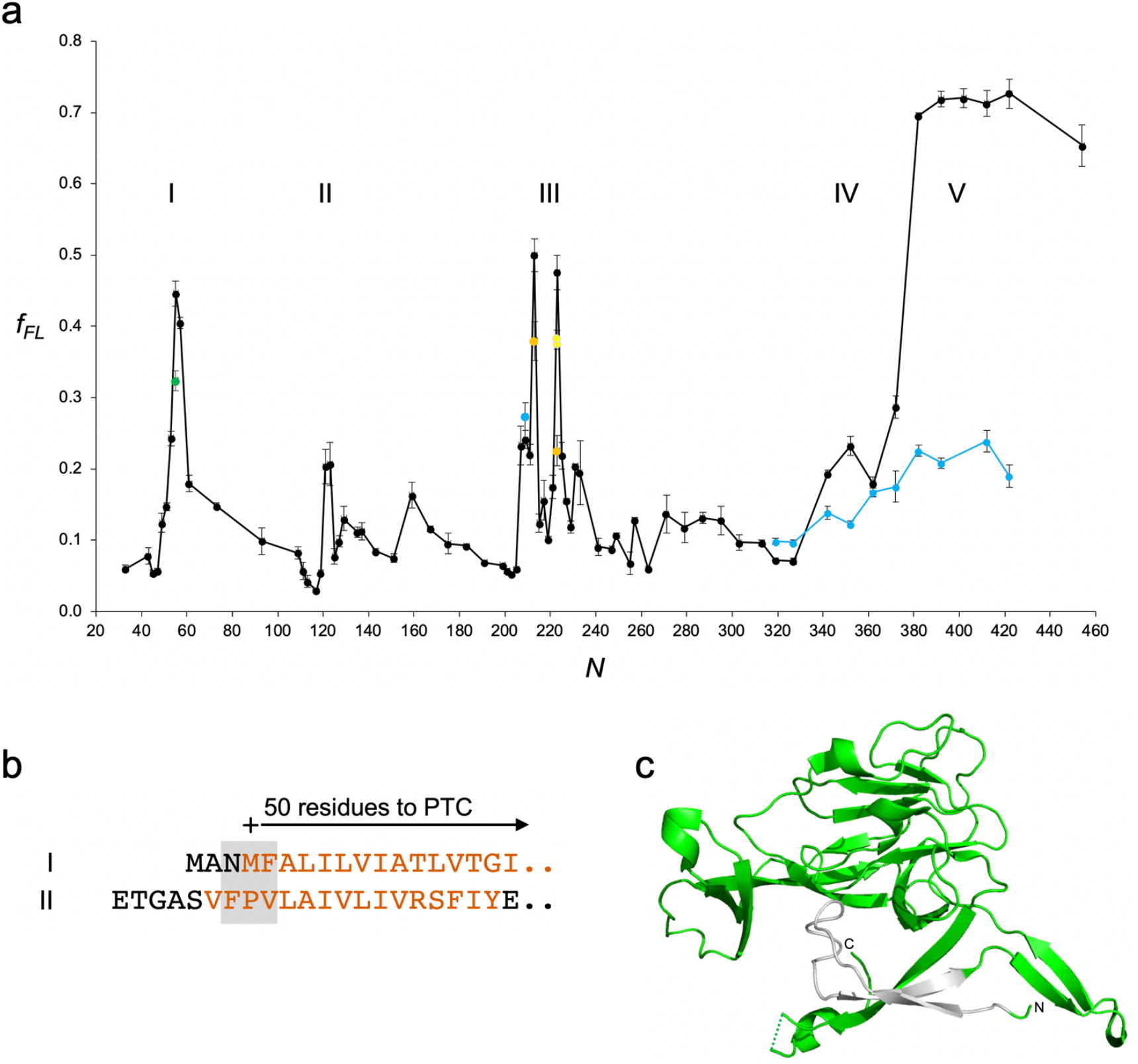
(a) FPs for LepB (black) and LepB[Δ80-105] (light blue). Mutants LepB[F^5^,L^7^→R^5^,R^7^] (*N* = 55; green), LepB[<u>Y</u>SNVE<u>P</u>S<u>DF</u>→<u>A</u>SNVE<u>A</u>S<u>AA</u>] (*N* = 213; orange),_LepB[<u>DF</u>VQT<u>F</u>S<u>RR</u>NGG<u>E</u> → <u>AA</u>VQT<u>A</u>S<u>AA</u>NGG<u>A</u>] (*N* = 223; orange), LepB[C^171^→A^171^] (N = 223; yellow), and LepB[C^177^→A^177^] (*N* = 223; yellow) are also shown. Error bars indicate SEM values. Note that the LepB[Δ80-105] constructs are 26 residues shorter than the corresponding LepB constructs but are plotted with the same *N*-values as the latter to simplify comparison. (b) Sequences with *N* = 53 and *N* = 120 (corresponding to *N*_*start*_ for peaks I and II) aligned from the critical Pro residue at the end of the AP. The + sign indicates the position 50 residues from the PTC in the arrested form of the protein, and the grey box shows the approximate uncertainty (±1 residue) in the determination of *N*_*start*_. Hydrophobic TMH segments are shown in orange. (c) Structure of the periplasmic domain of LepB (residues 78-324; PDB 1B12) with the [Δ80-105] deletion in grey.

Peak II reaches half-maximal amplitude at *N* = 120 residues, a nascent-chain length at which the N-terminal end of the weakly hydrophobic TMH2 is 52 residues away from the PTC, Fig. 2b. Peak II thus represents the membrane insertion of TMH2. *N*_*start*_ values obtained with the SecM(*Ms*) AP are typically ~5 residues larger than those obtained with the weaker SecM(*Ec*) AP [33], and the two TMHs thus probably reach the SecYEG translocon when their N-termini are ~45 residues away from the PTC, as seen for other *E. coli* inner membrane proteins [46].

The main peak in the FP is peak V. It has a much higher amplitude than the other peaks, and is also wider. It reaches its half-maximal amplitude at *N*_*start*_ ≈ 377 residues. Full-length LepB is 324 residues long, hence the C-terminal end of LepB is ~53 residues away from the PTC at this point, suggesting that peak V may represent the folding of the periplasmic domain. If this indeed is the case, the peak should disappear if the folded state is destabilized by mutation. We therefore deleted residues 80-105 that include a mostly buried segment in the core of the folded protein, Fig. 2c. This deletion abolished peak V (light blue data points in Fig. 2a). We conclude that folding of the periplasmic domain of LepB generates peak V by exerting sufficient force on the rather strong SecM(*Ms*) AP to overcome the translational arrest.

To determine if all or only a part of the periplasmic domain is involved in the peak V folding transition, we replaced an increasing number of C-terminal residues in LepB with Gly-Ser (GS) repeats and measured *f*_*FL*_ at *N* = 382. As seen in Fig. 3a, mutants with 3-6 GS repeats had significantly increased *f*_*FL*_ values, possibly because a more flexible linker gives the periplasmic domain more freedom to find an optimal location for folding relative to the SecYEG translocon. In contrast, longer replacements (8-14 GS repeats) progressively reduced *f*_*FL*_ towards the value seen for the non-folding LepB[Δ80-105] deletion mutant. Thus, the final ~15 residues of the periplasmic domain do not contribute to the folding transition seen at *N* = 382. These residues form a disordered loop and a short α-helix in the folded structure that have been proposed to help anchor the C-terminal end of LepB to the periplasmic surface of the inner membrane [47], Fig. 3b. We conclude that the main cotranslational folding transition happens when the C-terminal end of the last β-strand in the periplasmic domain is ~70 residues away from the PTC, and does not involve the last ~15 residues of the protein. Interestingly, peak V is more than 70 residues wide, and does not return to baseline even with a ~130-residue linker attached to LepB. It is thus apparent that membrane-attached, folded LepB (but not the unfolded Δ[80-105] mutant) exerts significant force on the AP even when it some distance away from the SecYEG translocon.

**Figure 3.**
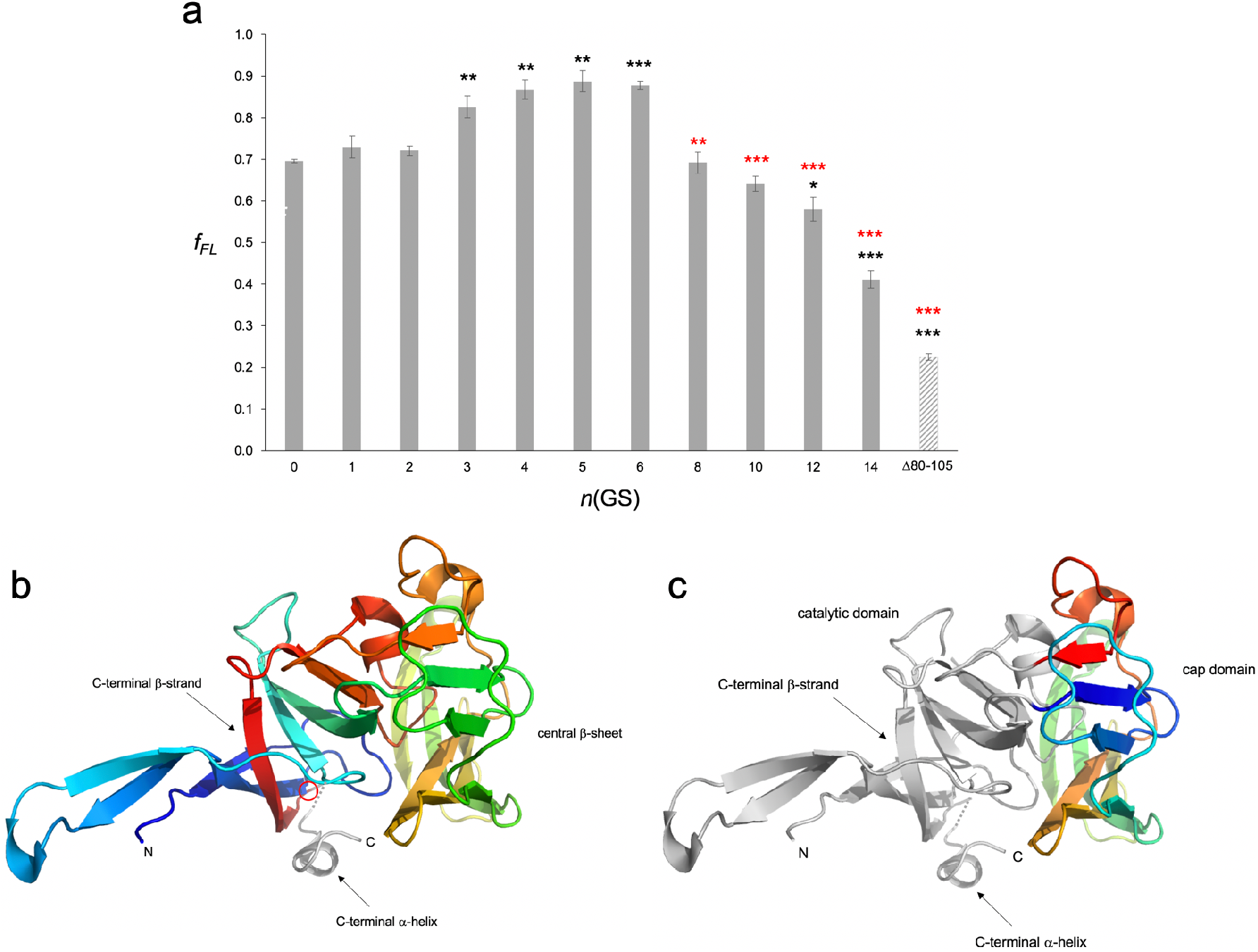
(a) *f*_*FL*_ values for LepB (*N* = 382) (GS)_n_ replacement mutants. *n* values are shown below the bars. The LepB[Δ80-105] (*N* = 382) construct is also included. *p*-values calculated by a two-sided t-test comparing the *f*_*FL*_ value for each construct to that for LepB (*N* = 382; black stars) and for LepB (*N* = 382_5GS; red stars) are indicated (*p* < 0.05, *; *p* < 0.01, **; *p* < 0.001, ***). (b) Structure of the periplasmic domain of LepB with the C-terminal disordered loop/α-helix in grey and residue 304 marked by a red circle. (c) Structure of the periplasmic domain of LepB with the cap domain (residues 153-263) in color.

Given that the distance between the PTC in the ribosome and the periplasmic end of the SecYEG translocon channel is ~160 Å [33], a distance that can be bridged by a largely extended nascent chain (~3.2 Å per residue) of ~50 residues length, we further conclude that the periplasmic domain has exited the translocon channel before it folds, and therefore that wildtype LepB folds post-translationally when not artificially tethered to the ribosome by a C-terminal linker.

Peaks III and IV are of lower amplitude and may signal the presence of folding intermediates in the P2 domain. Indeed, the deletion of residues 80-105 reduced the amplitude of peak IV (Fig. 2a, light blue curve), indicative of a folding intermediate. Peak IV reaches its half-maximal amplitude at *N* ≈ 330 residues and, assuming that the last ~70 residues are not part of this folding intermediate (as for the peak V folding transition), the folding intermediate would encompass approximately residues 1-260. Notably, residue 263 marks the end of the non-catalytic cap domain (also called domain II) that is found in only some of the LepB signal peptidases [48, 49], Fig. 3c, suggesting that peak IV may correspond to the folding of the cap domain.

Peak III, finally, is not reduced in amplitude by the residue 80-105 deletion, Fig. 2a (light blue data point at *N* = 209). The two narrow “spikes” at *N* = 213 and 223 seem to be caused by interactions between the nascent chain and the ribosome exit tunnel, as mutation of bulky and charged residues in the segments <u>Y</u>SNVE<u>P</u>S<u>DF</u> (located 21-29 residues from the PTC in the *N* = 213 construct) to <u>A</u>SNVE<u>A</u>S<u>AA</u> and <u>DF</u>VQT<u>F</u>S<u>RR</u>NGG<u>E</u> (located 20-32 residues from the PTC in the *N* = 223 construct) to <u>AA</u>VQT<u>A</u>S<u>AA</u>NGG<u>A</u>, markedly reduce the amplitude of the spikes (orange data points). We also considered the possibility that peak III may at least in part reflect the formation of the Cys^171^-Cys^177^ disulfide in the periplasmic domain (as we recently found for the periplasmic protein PhoA [50]); indeed, mutation of either Cys residue to Ala significantly reduces *f*_*FL*_ at *N* = 223 (*p* < 0.05, yellow data points). Because peak III does not seem to represent a major tertiary structure folding intermediate, we did not analyze it further.

## Discussion

Taken together with our previous analysis of the cotranslational folding of the periplasmic *E. coli* enzyme PhoA [50], the data presented here establish FPA as a generally applicable method to study the folding of periplasmic proteins and of periplasmic domains in inner membrane proteins. Applying FPA to the periplasmic domain of the inner membrane protease LepB, we find that the main folding transition does not involve the last ~15 residues of the protein, and takes place only when the domain has been fully translocated through the SecYEG translocon channel. This is not unexpected, given that the last β-strand in the periplasmic domain (residues 293-303) is required to connect the four N-terminal β-strands (residues 82-123) to the central and C-terminal β-sheet structures, Fig. 3b, and establishes that LepB normally folds post-translationally (when not tethered to the PTC via an added linker segment). The data further indicate that a less stable folding intermediate that includes the cap domain (residues 153-263) may form prior to the main folding transition.

Given that a ~50 residues long linker segment in an extended conformation should be able to reach from the PTC to the periplasmic end of the SecYEG channel, it is notable that the main folding transition in the periplasmic domain happens only at a linker length of ~70 residues. We speculate that this may be because the periplasmic domain is anchored to the inner membrane via the two N-terminal TMHs and by additional hydrophobic residues in the periplasmic domain itself, meaning that it will by necessity be located some distance away from the mouth of the SecYEG channel when it folds, as illustrated in Fig. 4.

**Figure 4.**
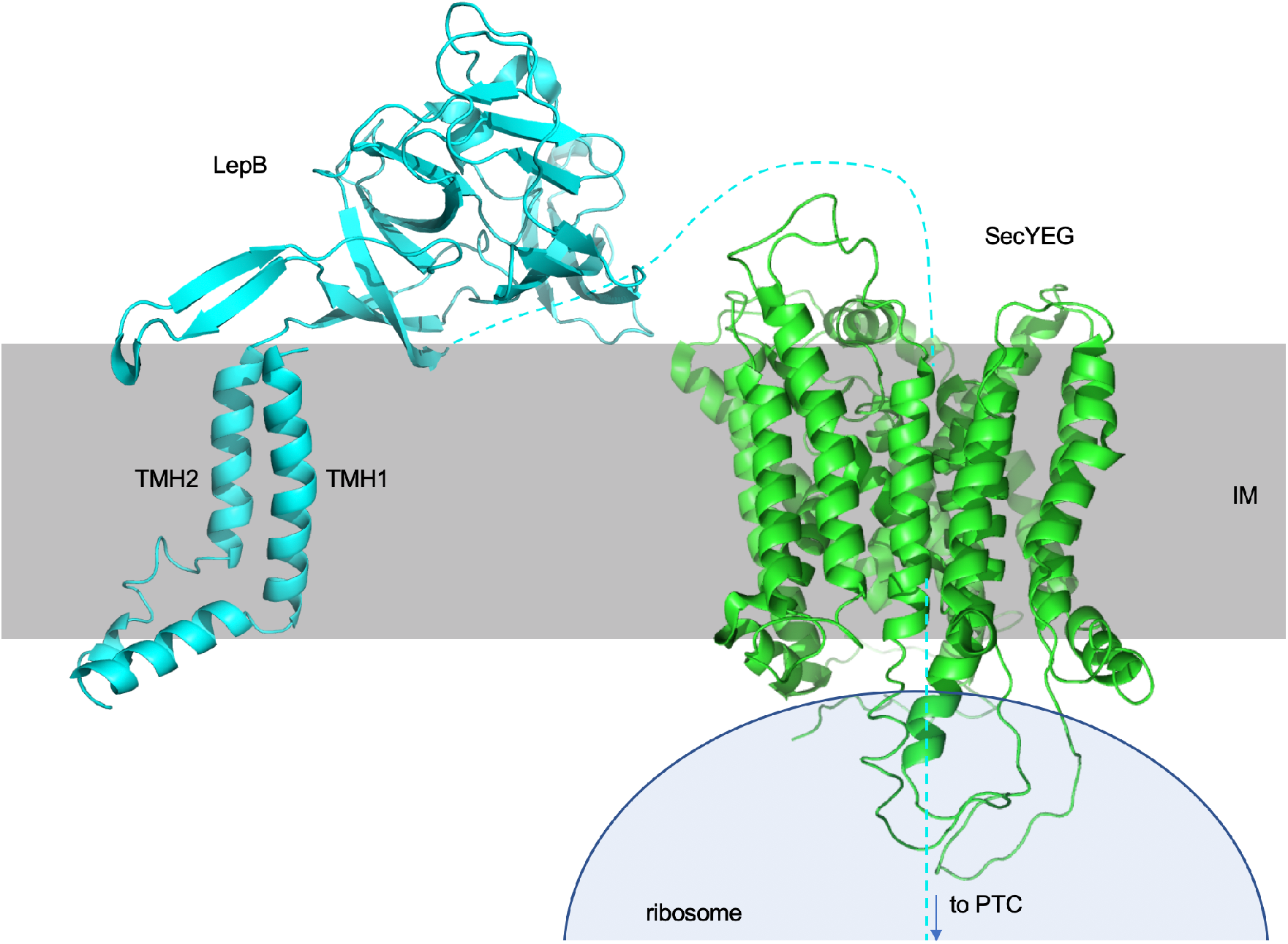
Model for LepB and SecYEG (PDB 5ABB) at the point of the main folding transition (*N* = 377). The folded periplasmic domain is shown in cartoon mode. The 70-residue linker connecting it to the PTC is dashed, and a part of the ribosome is sketched in light blue. The model for the TMH1-TMH2 membrane domain in LepB (residues 4-77) is the best-scoring prediction (Quality = 8) produced by the EVcouplings server [51] (https://v2.evcouplings.org), using LepB residues 1-93 as the seed.

Finally, it is quite remarkable that an AP located at the PTC, deep in the ribosome exit tunnel, can sense folding events taking place in the periplasm via a ~70 residues long linker that passes through both the ribosome and the SecYEG translocon. It will be interesting to test whether other events that take place in the periplasm, such as interactions between nascent polypeptides and periplasmic chaperones or steps in the assembly of outer membrane proteins, can be probed in a similar way.

## Materials and Methods

### Enzymes and chemicals

All enzymes used in this study were purchased from Thermo Fisher Scientific and New England Biolabs, with the exception of PfuUltra II Fusion HS DNA Polymerase that was procured from Stratagene, Sweden. Primers for site-directed mutagenesis, the partially overlapping inverse primers, and the primers used for Gibson Assembly^®^ were designed *in silico* and ordered from Eurofins Genomics. All gene fragments used for Gibson Assembly^®^ cloning were designed *in silico* and ordered using Invitrogen GeneArt Gene Synthesis service, Thermo Fisher Scientific. Plasmid isolation, PCR purification, gel extraction kits, and precast NuPAGE Bis-Tris polyacrylamide gels were from Thermo Fisher Scientific. L-[^35^S]-methionine was obtained from PerkinElmer. Mouse monoclonal antibody against the HA antigen was purchased from BioLegend. Protein-G agarose beads were manufactured by Roche. All other reagents were from Sigma-Aldrich.

### Cloning and Mutagenesis

Starting with the previously described pING plasmid carrying a variant of *lepB* gene with the *M. succiniciproducens* SecM(*Ms*) arrest peptide (AP) [33], the coding sequence for the 9-residue hemagglutinin (HA) tag and a 78-residue C-terminal tail were engineered upstream and downstream of the AP, respectively. In order to generate a force profile for LepB, the *lepB* sequence preceding the HA-tag was replaced with the wild-type *lepB* gene, amplified from the genomic DNA of MC1061 *E. coli* cells. The resulting *lepB-secM(Ms)* construct (*N* = 342), codes for the 324-residue LepB followed by a 96-residue chimeric domain that consists of the SecM(*Ms*) AP, an HA-tag, and a 78-residue C-terminal tail.

The pING plasmid containing the new *lepB-secM* construct (*N* = 342) was used as template to produce a series of constructs with progressively longer truncations of LepB, using partially overlapping inverse primers. The resulting *lepB-secM(Ms)* constructs of varying lengths are denoted by *N,* which is the number of residues from the N-terminus of LepB to the critical proline residue at C-terminal end of the AP.

During *in vivo* synthesis of the LepB-SecM(*Ms*) constructs, two different forms, the longer full-length (*FL*) form and the shorter arrested (*A*) form, are produced in amounts that depend on the force experienced by the AP. Controls for *FL* and *A* species were made using site-directed mutagenesis. *FL* controls have a non-functional AP that was generated by substituting the critical proline residue at the C-terminal end of the AP with an alanine. *A* controls have a stop codon at the C-terminal end of the AP to yield a product that is of equivalent size to the arrested species.

To follow translocation of the C-terminal periplasmic LepB domain across the inner membrane, the *lepB-secM(Ms)* construct *N* = 342 was extended by introducing additional HA-tags before the existing one through GeneArt Gibson Assembly^®^ Cloning. Gene fragments encoding multiple (2 to 9) HA-tags were designed *in silico* and ordered using GeneArt Gene Synthesis service from Thermo Fisher Scientific. Each additional 9-residue HA-tag was spaced by a methionine residue to extended the *lepB-secM(Ms)* constructs in steps of 10 residues, giving eight new constructs with lengths *N* = 352, 362, 372, 382, 392, 402, 412 and 422.

The *lepB-secM(Ms)* construct *N* = 422 was further extended by introducing a 32-residue PhoA segment before the HA-linkers through GeneArt Gibson Assembly^®^ Cloning. The gene fragment encoding the PhoA segment followed by nine HA-tags was designed *in silico* and ordered from GeneArt Gene Synthesis service, Thermo Fisher Scientific.

Non-folding control constructs were made for constructs *N* = 219, 319, 327, 342, 362, 382, 392, 412, and 422 by deleting residues 80-105 (FIYEPFQIPSGSMMPTLLIGDFILVE) in the LepB periplasmic domain by PCR using partially overlapping inverse primers.

In construct *N* = 382, C-terminal segments of LepB were replaced by stretches of glycine-serine (GS) repeats. The replacement was carried out using GeneArt Gibson Assembly^®^ Cloning. Gene fragments with successive replacement of LepB C-terminal sequence with codons for GS-repeats were designed *in silico* and ordered from GeneArt Gene Synthesis service, Thermo Fisher Scientific.

The hydrophobic residues Phe^5^ and Leu^7^ in LepB TMH1 were replaced with arginine (R) in construct *N* = 55 by site-directed mutagenesis using PfuUltra II Fusion HS DNA Polymerase (Stratagene, Sweden). Multiple alanine substitutions (underlined) were made in constructs *N* = 213 and 223 in residues 185-193 (<u>Y</u>SNVE<u>P</u>S<u>DF</u>) and 192-204 (<u>DF</u>VQT<u>F</u>S<u>RR</u>NGG<u>E</u>) using GeneArt Gibson Assembly^®^ Cloning. Two separate gene fragments with the underlined bulky and charged residues replaced with codons for alanine were designed *in silico* and ordered using GeneArt Gene Synthesis service, Thermo Fisher Scientific. Single alanine substitutions of Cys^171^ and Cys^177^ in construct *N* = 223 were made by site-directed mutagenesis using PfuUltra II Fusion HS DNA Polymerase (Stratagene, Sweden).

The PCR products were transformed into DH5α cells after DpnI treatment (1 hour at 37°C) and verified by agarose gel electrophoresis. Transformed cells were streaked on LB-agar plates containing 100 μg/ml carbenicillin (a semisynthetic ampicillin analog) for selection, and incubated at 37°C overnight. For plasmid isolation, single colonies were picked and cultured in LB supplemented with 100 μg/ml ampicillin and incubated in a rotary shaker for 16 hours at 37°C. Selectively replicated pING vectors containing the *lepB-secM(Ms)* constructs were subsequently purified from the cultured cells using GeneJET Plasmid Miniprep Kit from Thermo Scientific, according to the supplier’s specifications. Purified plasmid DNA was sent for sequencing at Eurofins Genomics to verify the sequence of the *lepB-secM(Ms)* constructs.

### In vivo pulse-labeling analysis

Competent *E. coli* MC1061 cells were transformed with the pING1 plasmids, and grown overnight at 37°C in M9 minimal medium supplemented with 19 amino acids (1 μg/ml, no Met), 100 μg/ml thiamine, 0.4% (w/v) fructose, 100 mg/ml ampicillin, 2 mM MgSO_4_, and 0.1mM CaCl_2_. Cells were diluted into fresh M9 medium to an OD_600_ of 0.1 and grown until an OD_600_ of 0.3 - 0.5.

LepB-SecM(*Ms*) protein expression was induced with 0.2% (w/v) arabinose and continued for 3 min at 37°C. Proteins were then radiolabeled with [^35^S]-methionine for 3 min at 37°C before the reaction was stopped by adding ice-cold trichloroacetic acid (TCA) to a final concentration of 10%. Samples were put on ice for 30 min and precipitates were spun down for 5 min at 20,000 g at 4°C in a tabletop centrifuge (Eppendorf). After one wash with ice-cold acetone, centrifugation was repeated and pellets were subsequently solubilized in Tris-SDS buffer (10 mM Tris-Cl pH 7.5, 2% (w/v) SDS) for 15 min at 37°C while shaking at 900 rpm. Samples were centrifuged for 5 min at 20,000 g to remove insoluble material. The supernatant containing solubilized protein was pre-cleared by incubation with Pansorbin^®^ in a buffer containing 50 mM Tris-HCl pH 8.0, 150 mM NaCl, 0.1 mM EDTA-KOH, 2% (v/v) Triton X-100. After 15 min incubation on ice, the suspension was centrifuged at 20,000 g and the supernatant incubated with Protein-G beads bound to Anti-HA.11 Epitope Tag Antibody (mouse) for immunoprecipitation. Binding and incubation was carried out overnight (16 hours) at 4°C on a roller.

After centrifugation for 1 min at 7,000 g, immunoprecipitates were first washed with 10 mM Tris-Cl pH 7.5, 150 mM NaCl, 2 mM EDTA, 0.2% (v/v) Triton X-100 and subsequently with 10 mM Tris-Cl pH 7.5. Samples were spun down again and pellets were solubilized in SDS sample buffer (67 mM Tris, 33% (w/v) SDS, 0.012% (w/v) bromophenol blue, 10 mM EDTA-KOH pH 8.0, 6.75% (v/v) glycerol, 100 mM DTT) for 10 min while shaking at 1,000 rpm. Samples were incubated with 0.25 mg/ml RNase I for 30 min at 37°C to hydrolyze the tRNA, and subsequently separated by SDS-PAGE. Gels were fixed in 30% (v/v) methanol and 10% (v/v) acetic acid and dried by using a Bio-Rad gel dryer model 583.

Radiolabeled proteins were detected by exposing dried gels to phosphorimaging plates, which were scanned in a Fuji FLA-3000 scanner. Band intensity profiles were obtained using the ImageGauge V4.23 software and quantified with our in-house software EasyQuant to determine the fraction full-length protein, *f*_*FL*_ = *I*_*FL*_/(*I*_*FL*_+*I*_*A*_), where *I*_*A*_, *I*_*FL*_ are the intensities of the *A* and *FL* bands, respectively. Data was collected from three to six independent biological replicates (see Supplementary Data), and averages and standard errors of the mean (SEM) were calculated. A two-sided t-test was used to calculate statistical significance when comparing *f*_*FL*_ values for different constructs.

## Abbreviations

PCR: polymerase chain reaction
TCA: trichloroacetic acid
SDS-PAGE: sodium dodecylsulphate polyacrylamide gel electrophoresis
PTC: peptidyl transferase center

## Author contributions

Sandhu: Formal Analysis, Investigation, Methodology, Writing- Review and editing; Hedman: Conceptualization, Formal Analysis, Methodology; Cymer: Conceptualization, Formal Analysis, Methodology, Writing-Review; Ismail: Conceptualization, Formal Analysis, Methodology, Writing- Review; Kudva: Conceptualization, Formal Analysis, Investigation, Methodology, Writing - Review and editing; von Heijne: Conceptualization, Formal Analysis, Funding acquisition, Project administration, Methodology, Resources, Supervision, Validation, Visualization, Writing - Original Draft, Writing - Review and editing

## Acknowledgements

This work was supported by grants from the Knut and Alice Wallenberg Foundation (2017.0323), the Novo Nordisk Fund (NNF18OC0032828), and the Swedish Research Council (621-2014-3713) to GvH.

## References

[1] Rapoport TA. Protein translocation across the eukaryotic endoplasmic reticulum and bacterial plasma membranes. Nature. 2007 450:663–669.

[2] Kadokura H, Beckwith J. Detecting folding intermediates of a protein as it passes through the bacterial translocation channel. Cell. 2009 138:1164–1173.

[3] Chen W, Helenius J, Braakman I, Helenius A. Cotranslational folding and calnexin binding during glycoprotein synthesis. Proc Natl Acad Sci U S A. 1995 92:6229–6233.

[4] Bergman LW, Kuehl WM. Formation of an intrachain disulfide bond on nascent immunoglobulin light chains. J Biol Chem. 1979 254:8869–8876.

[5] Kowarik M, Kung S, Martoglio B, Helenius A. Protein folding during cotranslational translocation in the endoplasmic reticulum. Mol Cell. 2002 10:769–778.

[6] Holtkamp W, Kokic G, Jäger M, Mittelstaet J, Komar AA, Rodnina MV. Cotranslational protein folding on the ribosome monitored in real time. Science. 2015 350:1104–1107.

[7] Liutkute M, Maiti M, Samatova E, Enderlein J, Rodnina MV. Gradual compaction of the nascent peptide during cotranslational folding on the ribosome. eLife. 2020 9.

[8] Frydman J, Erdjument-Bromage H, Tempst P, Hartl FU. Co-translational domain folding as the structural basis for the rapid de novo folding of firefly luciferase. Nat Struct Biol. 1999 6:697–705.

[9] Kelkar DA, Khushoo A, Yang Z, Skach WR. Kinetic analysis of ribosome-bound fluorescent proteins reveals an early, stable, cotranslational folding intermediate. J Biol Chem. 2012 287:2568–2578.

[10] Nicola AV, Chen W, Helenius A. Co-translational folding of an alphavirus capsid protein in the cytosol of living cells. Nat Cell Biol. 1999 1:341–345.

[11] Zhang G, Hubalewska M, Ignatova Z. Transient ribosomal attenuation coordinates protein synthesis and co-translational folding. Nat Struct Mol Biol. 2009 16:274–280.

[12] Samelson AJ, Bolin E, Costello SM, Sharma AK, O’Brien EP, Marqusee S. Kinetic and structural comparison of a protein’s cotranslational folding and refolding pathways. Sci Adv. 2018 4:eaas9098.

[13] Farias-Rico JA, Ruud Selin F, Myronidi I, Frühauf M, von Heijne G. Effects of protein size, thermodynamic stability, and net charge on cotranslational folding on the ribosome. Proc Natl Acad Sci U S A. 2018 115:E9280–E9287.

[14] Cassaignau AM, Launay HM, Karyadi ME, Wang X, Waudby CA, Deckert A, Robertson AL, Christodoulou J, Cabrita LD. A strategy for co-translational folding studies of ribosome-bound nascent chain complexes using NMR spectroscopy. Nature protocols. 2016 11:1492–1507.

[15] Deckert A, Waudby CA, Wlodarski T, Wentink AS, Wang X, Kirkpatrick JP, Paton JF, Camilloni C, Kukic P, Dobson CM, Vendruscolo M, Cabrita LD, Christodoulou J. Structural characterization of the interaction of alpha-synuclein nascent chains with the ribosomal surface and trigger factor. Proc Natl Acad Sci U S A. 2016 113:5012–5017.

[16] Eichmann C, Preissler S, Riek R, Deuerling E. Cotranslational structure acquisition of nascent polypeptides monitored by NMR spectroscopy. Proc Natl Acad Sci U S A. 2010 107:9111–9116.

[17] Waudby CA, Wlodarski T, Karyadi ME, Cassaignau AME, Chan SHS, Wentink AS, Schmidt-Engler JM, Camilloni C, Vendruscolo M, Cabrita LD, Christodoulou J. Systematic mapping of free energy landscapes of a growing filamin domain during biosynthesis. Proc Natl Acad Sci U S A. 2018 115:9744–9749.

[18] Cabrita LD, Cassaignau AM, Launay HM, Waudby CA, Wlodarski T, Camilloni C, Karyadi ME, Robertson AL, Wang X, Wentink AS, Goodsell LS, Woolhead CA, Vendruscolo M, Dobson CM, Christodoulou J. A structural ensemble of a ribosome-nascent chain complex during cotranslational protein folding. Nat Struct Mol Biol. 2016 23:278–285.

[19] Cabrita LD, Hsu ST, Launay H, Dobson CM, Christodoulou J. Probing ribosome-nascent chain complexes produced in vivo by NMR spectroscopy. Proc Natl Acad Sci U S A. 2009 106:22239–22244.

[20] Nilsson OB, Hedman R, Marino J, Wickles S, Bischoff L, Johansson M, Muller-Lucks A, Trovato F, Puglisi JD, O’Brien EP, Beckmann R, von Heijne G. Cotranslational protein folding inside the ribosome exit tunnel. Cell Rep. 2015 12:1533–1540.

[21] Kudva R, Tian P, Pardo-Avila F, Carroni M, Best RB, Bernstein HD, von Heijne G. The shape of the bacterial ribosome exit tunnel affects cotranslational protein folding. eLife. 2018 7:e36326.

[22] Tian P, Steward A, Kudva R, Su T, Shilling PJ, Nickson AA, Hollins JJ, Beckmann R, von Heijne G, Clarke J, Best RB. Folding pathway of an Ig domain is conserved on and off the ribosome. Proc Natl Acad Sci U S A. 2018 115:E11284–E11293.

[23] Katranidis A, Grange W, Schlesinger R, Choli-Papadopoulou T, Bruggemann D, Hegner M, Buldt G. Force measurements of the disruption of the nascent polypeptide chain from the ribosome by optical tweezers. FEBS Lett. 2011 585:1859–1863.

[24] Wruck F, Katranidis A, Nierhaus KH, Buldt G, Hegner M. Translation and folding of single proteins in real time. Proc Natl Acad Sci U S A. 2017 114:E4399–E4407.

[25] Kaiser CM, Goldman DH, Chodera JD, Tinoco IJr.,, Bustamante C. The ribosome modulates nascent protein folding. Science. 2011 334:1723–1727.

[26] Goldman DH, Kaiser CM, Milin A, Righini M, Tinoco I, Bustamante C. Mechanical force releases nascent chain-mediated ribosome arrest *in vitro* and *in vivo*. Science. 2015 348:457–460.

[27] Nilsson OB, Müller-Lucks A, Kramer G, Bukau B, von Heijne G. Trigger factor reduces the force exerted on the nascent chain by a cotranslationally folding protein. J Mol Biol. 2016 428:1356–1364.

[28] Nilsson OB, Nickson AA, Hollins JJ, Wickles S, Steward A, Beckmann R, von Heijne G, Clarke J. Cotranslational folding of spectrin domains via partially structured states. Nat Struct Mol Biol. 2017 24:221–225.

[29] Farias-Rico JA, Goetz SK, Marino J, von Heijne G. Mutational analysis of protein folding inside the ribosome exit tunnel. FEBS Lett. 2017 591:155–163.

[30] Kemp G, Kudva R, de la Rosa A, von Heijne G. Force-Profile Analysis of the Cotranslational Folding of HemK and Filamin Domains: Comparison of Biochemical and Biophysical Folding Assays. J Mol Biol. 2019 431:1308–1314.

[31] Jensen MK, Samelson AJ, Steward A, Clarke J, Marqusee S. The folding and unfolding behavior of ribonuclease H on the ribosome. J Biol Chem. 2020 295:11410–11417.

[32] Ito K, Chiba S. Arrest peptides: cis-acting modulators of translation. Annu Rev Biochem. 2013 82:171–202.

[33] Ismail N, Hedman R, Schiller N, von Heijne G. A biphasic pulling force acts on transmembrane helices during translocon-mediated membrane integration. Nature Struct Molec Biol. 2012 19:1018–1022.

[34] Kemp G, Nilsson OB, Tian P, Best RB, von Heijne G. Cotranslational folding cooperativity of contiguous domains of α-spectrin. Proc Natl Acad Sci U S A. 2020 117:14119–14126.

[35] Shanmuganathan V, Schiller N, Magoulopoulou A, Cheng J, Braunger K, Cymer F, Berninghausen O, Beatrix B, Kohno K, von Heijne G, Beckmann R. Structural and mutational analysis of the ribosome-arresting human XBP1u. eLife. 2019 8.

[36] Cymer F, Hedman R, Ismail N, von Heijne G. Exploration of the arrest peptide sequence space reveals arrest-enhanced variants. J Biol Chem. 2015 290:10208–10215.

[37] Wolfe PB, Wickner W, Goodman JM. Sequence of the leader peptidase gene of *Escherichia coli* and the orientation of leader peptidase in the bacterial envelope. J Biol Chem. 1983 258:12073–12080.

[38] Paetzel M, Dalbey RE, Strynadka NCJ. Crystal structure of a bacterial signal peptidase apoenzyme - Implications for signal peptide binding and the Ser-Lys dyad mechanism. Journal of Biological Chemistry. 2002 277:9512–9519.

[39] de Gier J-WL, Mansournia P, Valent Q, Phillips GJ, Luirink J, von Heijne G. Assembly of a cytoplasmic membrane protein in *Escherichia coli* is dependent on the signal recognition particle. FEBS Lett. 1996 399:307–309.

[40] Valent QA, de Gier JWL, von Heijne G, Kendall DA, ten Hagen-Jongman CM, Oudega B, Luirink J. Nascent membrane and presecretory proteins synthesized in *Escherichia coli* associate with signal recognition particle and trigger factor. Molecular microbiology. 1997 25:53–64.

[41] Schibich D, Gloge F, Pöhner I, Björkholm P, Wade RC, von Heijne G, Bukau B, Kramer G. Global profiling of SRP interaction with nascent polypeptides. Nature. 2016 536:219–223.

[42] Wolfe PB, Rice M, Wickner W. Effects of two *sec* genes on protein assembly into the plasma membrane of *Escherichia coli*. J Biol Chem. 1985 260:1836–1841.

[43] Houben EN, Zarivach R, Oudega B, Luirink J. Early encounters of a nascent membrane protein: specificity and timing of contacts inside and outside the ribosome. J Cell Biol. 2005 170:27–35.

[44] Houben EN, ten Hagen-Jongman CM, Brunner J, Oudega B, Luirink J. The two membrane segments of leader peptidase partition one by one into the lipid bilayer via a Sec/YidC interface. EMBO Rep. 2004 5:970–975.

[45] Mercier E, Wintermeyer W, Rodnina MV. Co-translational insertion and topogenesis of bacterial membrane proteins monitored in real time. EMBO J. 2020 e104054.

[46] Nicolaus F, Metola A, Mermans D, Liljenström A, Krč A, Abdullahi SM, Zimmer M, Miller TF, von Heijne G. Residue-by-residue analysis of cotranslational membrane protein integration *in vivo*. BioRxiv. 2020 2020.2009.2027.315283.

[47] Paetzel M, Dalbey RE, Strynadka NCJ. Crystal structure of a bacterial signal peptidase in complex with a β-lactam inhibitor. Nature. 1998 396:186–190.

[48] Paetzel M, Karla A, Strynadka NC, Dalbey RE. Signal peptidases. Chem Rev. 2002 102:4549–4580.

[49] Young PG, Proft T, Harris PW, Brimble MA, Baker EN. Structure and activity of Streptococcus pyogenes SipA: a signal peptidase-like protein essential for pilus polymerisation. PLoS One. 2014 9:e99135.

[50] Elfageih R, Karyolaimos A, Kemp G, de Gier JW, von Heijne G, Kudva R. Cotranslational folding of alkaline phosphatase in the periplasm of Escherichia coli. Protein Sci. 2020 29:2028–2037.

[51] Hopf TA, Green AG, Schubert B, Mersmann S, Scharfe CPI, Ingraham JB, Toth-Petroczy A, Brock K, Riesselman AJ, Palmedo P, Kang C, Sheridan R, Draizen EJ, Dallago C, Sander C, Marks DS. The EVcouplings Python framework for coevolutionary sequence analysis. Bioinformatics. 2019 35:1582–1584.

